# Transcriptome profiling reveals differential expression of virulence genes in *Xylella fastidiosa* under nutrient-rich and xylem-like conditions

**DOI:** 10.1101/2025.08.15.669762

**Authors:** Paulo M. Pierry, Oseias R. Feitosa-Junior, Joaquim Martins-Junior, Deibs Barbosa, Aline M. da Silva, Paulo A. Zaini

## Abstract

*Xylella fastidiosa* is a xylem-limited, insect-vectored pathogen. To capture the metabolic changes that underlie its adaptation to both nutrient-rich and xylem-like environments, we performed transcriptome analyses on two reference strains, Temecula1 and 9a5c, sampled at early and late exponential phases in a complex laboratory medium (PWG) and a defined, xylem-mimicking medium (PIM6). Across eight conditions (24 transcriptomes), over 90% of annotated coding sequences were expressed, although 40–80% of transcripts exhibited low abundance (TPM < 100). In contrast, a core set of highly abundant transcripts was seen in both strains and media, including the non-coding RNAs 6S and RNase P, as well as multiple colicin- and microcin-like toxins, proteases, lipases, and stress-response factors. Comprehensive pathway reconstruction confirmed the activity of all essential metabolic pathways, while revealing conditional enrichment of stress-related and defense modules under nutrient limitation. We annotated 5′ and 3′ untranslated regions for thousands of transcripts and defined over 900 operons across both genomes. Differential expression analysis demonstrated that PIM6 medium confers a far greater transcriptional reprogramming between early and late phases than PWG. Strain-to-strain comparisons revealed that up to 12.6% of orthologous genes were differentially regulated, emphasizing divergent regulatory networks even under identical growth conditions. This multi-dimensional transcriptome atlas not only refines *X. fastidiosa* genome annotation but also pinpoints candidate regulators, non-coding RNAs, and operon variants for functional validation. By illuminating the dynamic gene expression landscapes that enable xylem colonization and environmental resilience, our study lays a foundation for targeted disruption of key pathways in this important plant pathogen.

## INTRODUCTION

Transcriptome profiling via RNA sequencing (RNA-Seq) has become a go-to approach for elucidating how plant-pathogenic bacteria alter their gene expression in response to varying nutrient conditions. Wide changes in gene expression have been reported for several phytopathogenic bacteria, affecting type III secretion system components and effectors (Liu et al., 2021), quorum sensing (Kang et al., 2022; Sibanda et al., 2018), phenotype conversion (Nakahara et al., 2021), and nutrient transport (Liu et al., 2013), among many other traits. In this context, *Xylella fastidiosa* is an ideal model to be investigated, since its life cycle alternates between two entirely contrasting niches – that of the plant xylem and insect mouthparts (Chatterjee et al., 2008). Despite studies that investigated gene expression in different media (Amoia et al., 2025; Ciraulo et al., 2010; Pashalidis et al., 2005), systematic comparisons of growth in these two conditions remain limited.

The draft genomes of two reference strains—9a5c, obtained from a symptomatic sweet orange in Brazil, and Temecula1, isolated from an infected grapevine in California—were released in the early 2000s (Simpson et al., 2000; Van Sluys et al., 2003) paving the way for the first transcriptomic investigations. Early DNA microarray studies with these strains demonstrated that cells freshly extracted from plant xylem vessels exhibit a transcriptional profile enriched in cell-wall-degrading enzymes, adhesion factors, and detoxification systems relative to cells grown *in vitro*, suggesting that passage through the host primes *X. fastidiosa* for persistent colonization (De Souza et al., 2003). Subsequent comparisons of biofilm versus planktonic cultures revealed a reciprocal regulation of motility genes and exopolysaccharide biosynthesis pathways, highlighting the dynamic balance between dispersal and surface attachment that underlies both virulence and transmission (Silva et al., 2011).

Several other sequencing- and microarray-based studies (Da Silva Neto et al., 2010, 2007; Dourado et al., 2015; Fogaça et al., 2010; Muranaka et al., 2013; Nunes et al., 2003; Travensolo et al., 2009; Voegel et al., 2013; Wang et al., 2012; Zaini et al., 2008) have greatly advanced our understanding of *X. fastidiosa* biology. However, they inherently lacked the resolution to pinpoint transcriptional start sites, delineate untranslated regions, or accurately reconstruct operon structures. RNA-Seq overcomes these limitations by providing single-base resolution of transcript boundaries and quantitative measurements across a vast dynamic range (Sorek and Cossart, 2010). To date, three studies have applied RNA-Seq directly to *X. fastidiosa* under controlled *in vitro* conditions. The first novel small RNAs and operon variants linked to biofilm maturation in *X. fastidiosa* Temecula1 were revealed under calcium-supplemented conditions (Parker et al., 2016). A follow-up study using microfluidic chambers that mimic xylem flow further demonstrated calcium-dependent regulation of type IV pili, recombination, and signaling pathways (Chen and De La Fuente, 2020). More recently, co-cultivation with the endophyte *Methylobacterium mesophilicum* revealed transcriptional reprogramming of metabolic and transport pathways in response to nutrient competition (Dourado et al., 2023). However, most RNA-Seq efforts focused on plant responses to *Xylella fastidiosa* infection—profiling changes in *Arabidopsis thaliana* (Rogers, 2012), *Citrus reticulata* (Pereira et al., 2020), *Olea europaea* (olive) (Giampetruzzi et al., 2016), *Vitis vinifera* (grapevine) (Zaini et al., 2018), and even natural infections in olive through pilot dual RNA-Seq (Amoia et al., 2025), while the bacterial transcriptome continues to be relatively underexplored.

Here, we present the first comprehensive RNA-Seq analysis of *X. fastidiosa* strains 9a5c and Temecula1, sampled during exponential growth in two contrasting media: a complex medium (PWG) routinely used for laboratory cultivation, and a defined medium (PIM6) formulated to mimic the nutrient composition of xylem sap. By integrating high-resolution transcript mapping with global expression profiling, we aimed to (i) annotate untranslated regions and operon architecture across the genome, (ii) compare strain-specific and media-dependent transcriptional programs, and (iii) identify regulatory pathways enriched under xylem-like conditions. These data improve *X. fastidiosa* genome annotation, uncover virulence-associated regulatory profiles, and highlight molecular strategies this pathogen employs to persist in xylem vessels and adapt to contrasting environments, informing targeted disease management strategies.

## MATERIAL AND METHODS

### *Xylella fastidiosa* strains and growth conditions

*Xylella fastidiosa* strains 9a5c (Li et al., 1999) and Temecula1 (Van Sluys et al., 2003) were grown on PWG medium (PW supplemented with 0.5% glucose and 0.4% glutamine; (Davis et al., 1981)) or on PWG agar (1.5% agar) at 28 °C. Cultures were refreshed from −80 °C glycerol stocks after 20 passages. For liquid cultures, cells from 7–10-day PWG-agar plates were inoculated into 50–100 mL PWG or PIM6 medium (HEPES 10 mM pH 6.5, MgSO₄ 1 mM, CaCl₂ 3 mM, D-glucose 0.1 mM, sodium tartrate 0.1 mM, sodium malate 0.2 mM, sodium citrate 1 mM, K₂HPO₄ 0.05 g L⁻¹, KH₂PO₄ 0.03 g L⁻¹, L-glutamine 5 mM, ATCC13061 micronutrients 0.025%, soytone 0.02%, tryptone 0.04%) and incubated at 28 °C, 170 rpm (up to 15 days for PWG, 3 days for PIM6). Growth curves were obtained in 10 mL medium in 50 mL tubes inoculated at OD_600nm_ = 0.05 and measured at days 1, 3, 7, 10, 15, and 21 after vortexing to detach cells.

### RNA extraction and quality control

Cultures were initiated at a concentration of OD_600nm_ = 0.05 (PWG) or 0.25 (PIM6; after 24 h acclimation at OD_600nm_ = 0.3). Three biological replicates were collected per condition. To ensure comprehensive transcriptome coverage, cells were collected from the entire culture flask, including both the planktonic phase and those adhered to the glass surface. For this, the culture medium was thoroughly pipetted and flushed along the flask walls to dislodge adherent cells prior to harvesting. PIM6 samples were treated with LifeGuard reagent (Mobio) for RNA stabilization. Total RNA was extracted using Trizol (Invitrogen), followed by column purification (PureLink RNA Mini Kit, Ambion). DNase treatment (Illustra RNASpin, GE Healthcare) was applied with increased enzyme volume and incubation (50 min, RT), followed by re-purification. RNA integrity (RIN > 7.5) was assessed by BioAnalyzer (Agilent) and quantified by NanoDrop. Absence of DNA contamination was verified by PCR using CVC-1/272-2-int primers (Pooler and Hartung, 1995).

### Library preparation and sequencing

rRNA was depleted (Ribo-Zero Magnetic Kit, Illumina) from 1–5 µg total RNA. Residual RNA was precipitated and directly resuspended in TruSeq Elute, Prime, Fragment Mix. Libraries were prepared using TruSeq RNA Sample Prep Kit v2 (Illumina) and quality-checked on a BioAnalyzer. Quantification was performed by qPCR (KAPA Library Quantification Kit). Sequencing was done on MiSeq (Illumina) with 2 × 250 bp paired-end reads. A total of 24 libraries were sequenced in five runs (6–10 pM loading), yielding >25 M reads/run, with 75–82% high-quality reads and 70–90% mapping to reference genomes (Table S1). Coverage exceeded 300× for all samples.

### Transcriptome processing and differential expression

Read quality was assessed with FastQC (Andrews, S., 2010). Although quality trimming (>Q30) was initially performed using CLC Genomics Workbench to remove low-quality bases, it was observed that this stringent criterion led to excessive sequence shortening, potentially compromising genome mapping. Therefore, untrimmed paired-end reads were used for downstream differential expression analysis. Mapping to reference genomes was performed using CLC Genomics Workbench (v6.5) with RNA-Seq analysis, generating FPKM and raw count tables. FPKM values were converted to TPM for within-sample normalization. Coverage was computed as C = (L × N)/G, where L is the read length, N is the number of reads, and G is the genome size. Differential expression was analyzed with DESeq2 (Love, Huber, and Anders 2014) using FDR-adjusted *p* ≤ 0.05 and |log₂FC| > 1. Operon prediction was refined using *Rockhopper2* (Tjaden, 2015) and compared to *DOOR2* (Dam et al., 2007; Mao et al., 2009). Functional analysis employed *MinPath* (Ye and Doak, 2009); genome comparisons used IMG/ER (Markowitz et al., 2012), and sequence similarity searches were performed with BLAST (Altschul et al., 1990).

### Trend and global analyses

Common genes between the strains were identified from locus-tag correspondences. Expression trends were analyzed via two strategies: (i) top 100 expressed genes and (ii) 73 virulence-associated genes (Uceda-Campos et al., 2022). ANOVA with FDR correction and Tukey’s HSD was used for significance testing. Visualizations were generated using Python (pandas, seaborn, matplotlib). Global variation was explored by (i) PCoA, PERMANOVA, and PERMDISP (scikit-bio), (ii) expression-distribution bubble plots, and (iii) UpSet plots (UpSetPlot).

### GO enrichment and clustering analyses

DEGs were functionally classified by GO terms and analyzed for enrichment using BayGO (Vêncio et al., 2006) with G-test and FDR correction. Significant Biological Process terms (*p* ≤ 0.05) were visualized as comparative bar plots. Hierarchical clustering of log_2_FC values was performed in seaborn (clustermap), applying a custom diverging colormap.

## RESULTS

### Growth dynamics and adhesion phenotypes

To determine the appropriate time points for RNA sampling during early and late exponential growth, strains 9a5c and Temecula1 were cultured for 21 days in rich (PWG) and minimal (PIM6) media. As expected, both strains reached higher optical densities (OD_600nm_) in PWG than in PIM6. In the minimal medium, stationary phase was reached earlier (∼3 days) (Figure 1A–B). In the rich PWG medium, strain 9a5c sustained exponential growth for a longer period (∼10 days) than Temecula1 (7 days). Both strains adhered to glass surfaces regardless of the culture medium — a common phenotype of *X. fastidiosa in vitro* (Feitosa-Junior et al., 2022; Shi et al., 2009). However, strain 9a5c exhibited significantly greater surface adhesion than Temecula1, forming a thicker biofilm at the air–liquid interface (Figure 1C–J).

**Figure 1.**
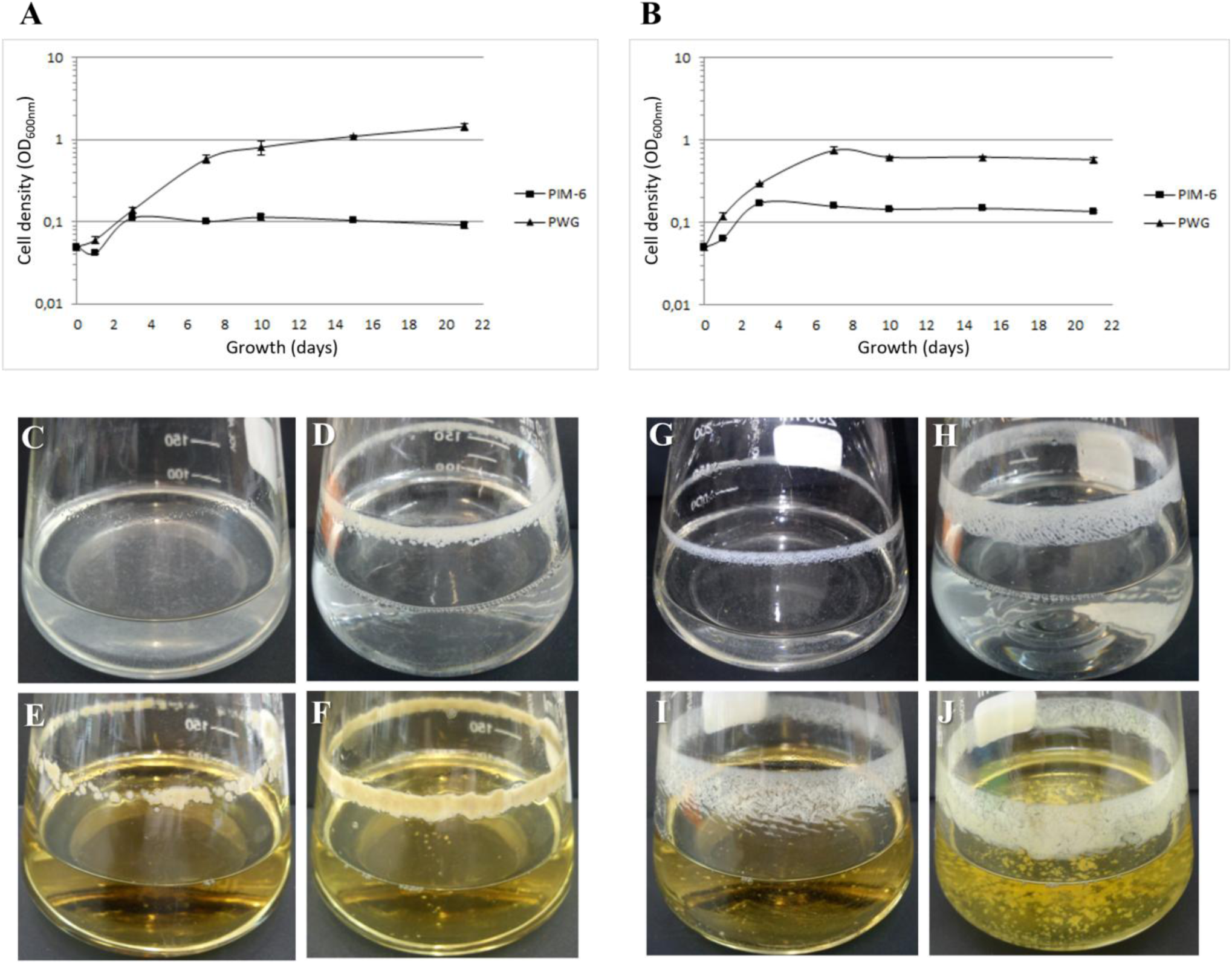
Growth curves of strains 9a5c and Temecula1 cultured in PIM6 and PWG media and their respective phenotypes. **(A)** Cells of strains 9a5c **(B)** and Temecula1 were grown in 50 mL conical tubes with initial OD600nm = 0.05 in PWG (▲) and PIM6 (▪) media, at 28 °C and 170 rpm. Measurements of OD600nm were done at days 1, 3, 7, 10, 15, and 21. Vertical bars represent the standard deviation of the mean of three technical replicates. Images on the lower panels represent the *in vitro* culture of time periods chosen for RNA-Seq analysis, based on the above curves: **(C)** 9a5c-PIM6-early, **(D)** 9a5c-PIM6-late, **(E)** 9a5c-PWG-early, **(F)** 9a5c-PWG-late, **(G)** Tem1-PIM6-early, **(H)** Tem1-PIM6-late, **(I)** Tem1-PWG-early, **(J)** Tem1-PWG-late.

Cells were collected at 1 day (PIM6) and 3 days (PWG) for early exponential phase transcriptomes. For late exponential phase, samples were collected at 3 days in PIM6, and at 7 days (Temecula1) or 10 days (9a5c) in PWG.

### Transcriptome architecture: UTRs, operons & non-coding RNAs

Given the high quality and consistency of the replicate samples (Figure 2; Table S2), transcript sequences were used to define the 5′ and 3′ untranslated regions (UTRs) and to predict operon structures in both genomes.

**Figure 2.**
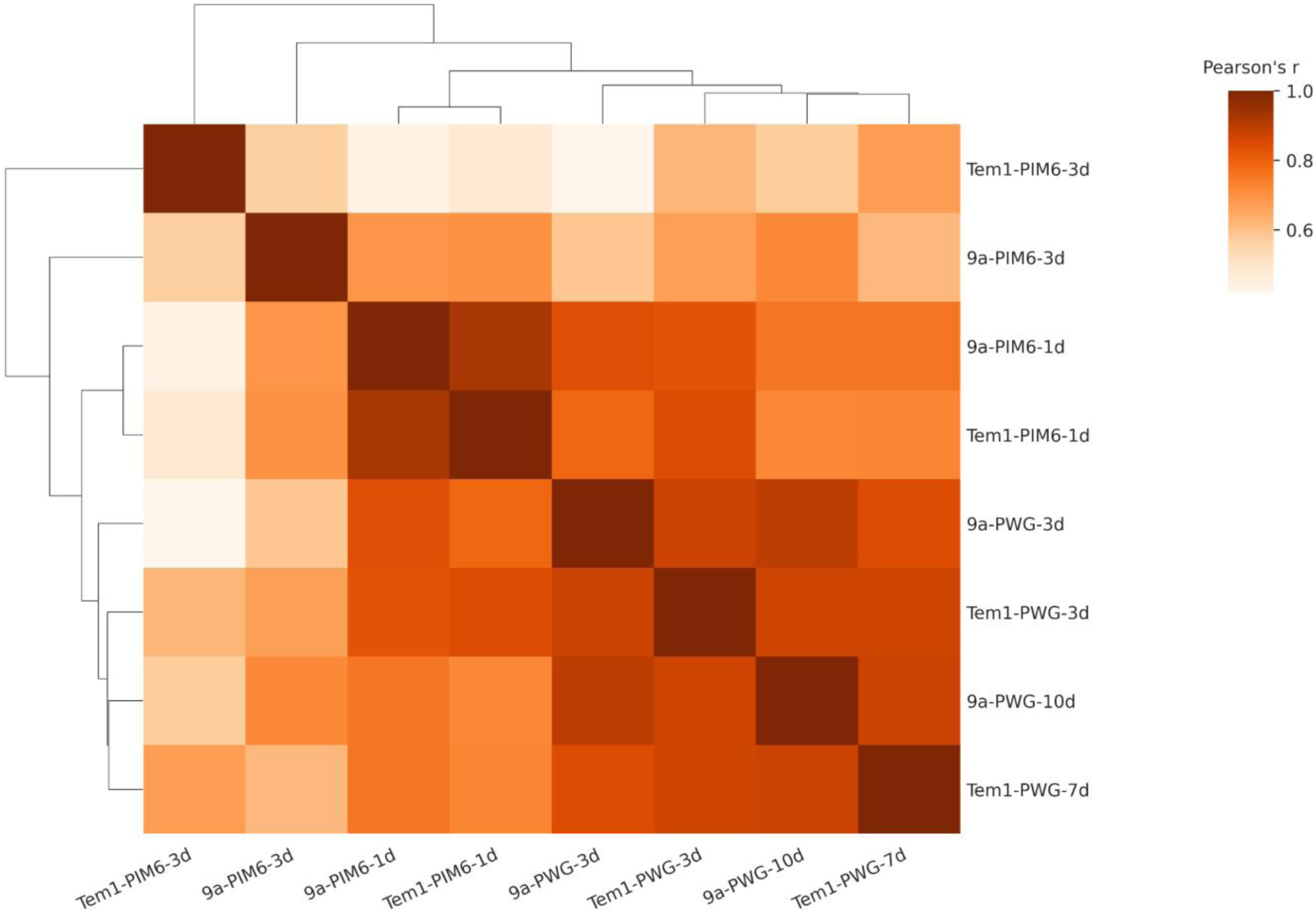
Hierarchical clustering of transcriptomic correlations among experimental conditions. The clustered heatmap shows pairwise Pearson correlation coefficients (TPM-based) across eight sample groups, varying by strain (9a5c or Temecula1), culture medium (PIM6 or PWG), and time point. Samples are hierarchically clustered using Euclidean distance and average linkage. Warmer tones indicate stronger positive correlations. The color bar reflects the Pearson correlation coefficient (r).

In strain 9a5c, we identified 1,204 5′UTRs (41.3% of genes) and 1,245 3′UTRs (46.3%). For the pXF51 plasmid, 31 5′UTRs (51.6%) and 25 3′UTRs (46.9%) were annotated. In comparison, Temecula1 showed 899 5′UTRs (38.3%) and 987 3′UTRs (46.5%). Supplementary tables S3 and S4 describe the coordinates of these UTR regions found in both strains.

Operon structures were also predicted based on the identification of polycistronic transcripts. A total of 544 operons were predicted in 9a5c and 386 in Temecula1 (Table 1). Most operons were predicted to contain only two genes, although larger operons with more than 10 genes were also described (Table 1). One of these large operons, spanning genes XF0305 to XF0318, includes genes involved in the oxidative phosphorylation pathway. Operon predictions from RNA-Seq data were compared with *in silico* annotations from the DOOR2 database (Dam *et al*., 2007; Mao *et al*., 2009). Table 1 shows a similar number of operons predicted for strains 9a5c and Temecula1, although some discrepancies in operon structure were observed.

**Table 1.** Prediction of operon structures of *X. fastidiosa* strains, using Rockhopper2 and the DOOR^2^ database. Columns indicate the number of genes in the operon.

| Strain | Method | Total | Number of genes per operon |  |  |  |  |  |  |  |  |
| --- | --- | --- | --- | --- | --- | --- | --- | --- | --- | --- | --- |
|  |  |  | 2 | 3 | 4 | 5 | 6 | 7 | 8 | 9 | ≥10 |
| 9a5c | RNA-Seq | 544 | 314 | 114 | 56 | 23 | 12 | 8 | 4 | 5 | 8 |
|  | DOOR<sup>2</sup> | 546 | 323 | 117 | 60 | 22 | 6 | 4 | 5 | 3 | 6 |
| 9a5c<br>pXF51 | RNA-Seq | 14 | 7 | 1 | 2 | 2 | 0 | 1 | 1 | 0 | 0 |
|  | DOOR<sup>2</sup> | 13 | 7 | 1 | 3 | 0 | 0 | 1 | 1 | 0 | 0 |
| Temecula1 | RNA-Seq | 386 | 209 | 88 | 46 | 19 | 11 | 4 | 2 | 2 | 5 |
|  | DOOR<sup>2</sup> | 400 | 233 | 84 | 42 | 22 | 7 | 4 | 2 | 1 | 5 |

Although strand-specific libraries were not used, all predicted operons contained genes in the same orientation, as manually annotated. Supplementary Tables S5 and S6 list all operons identified from RNA-Seq data.

In all transcriptomes, the most highly expressed transcript (based on TPM) corresponded to the non-coding RNA component of bacterial RNase P (class A). 6S RNA was also consistently abundant, ranking among the top ten transcripts in 19 of the 24 transcriptomes.

### Global expression profiles and plasmid-encoded traits

Normalized TPM values were used to compare the expression level of genes within a given transcriptome. Principal Coordinates Analysis (PCoA) on Bray–Curtis distances segregated samples by strain, medium, and growth phase (PERMANOVA p < 0.005; PERMDISP p < 0.01), with PC1 capturing 52.9 % of variance. This axis highlighted tighter clustering of PIM6 samples versus PWG, and greater dispersion in early-exponential than late-exponential cultures (Figure 3A). Genes in each transcriptome were ranked by TPM expression values and grouped into five categories: ultra-high (>10,000), high (1,000–10,000), moderate (100–1,000), low (0–100), and absent (0). Most genes showed low expression (TPM between 0 and 100). These normalized values show that, despite a subset with very high TPM values, most of the genes (44.9% to 70.4%) showed low expression in all transcriptomes analyzed (Figure 3B).

**Figure 3.**
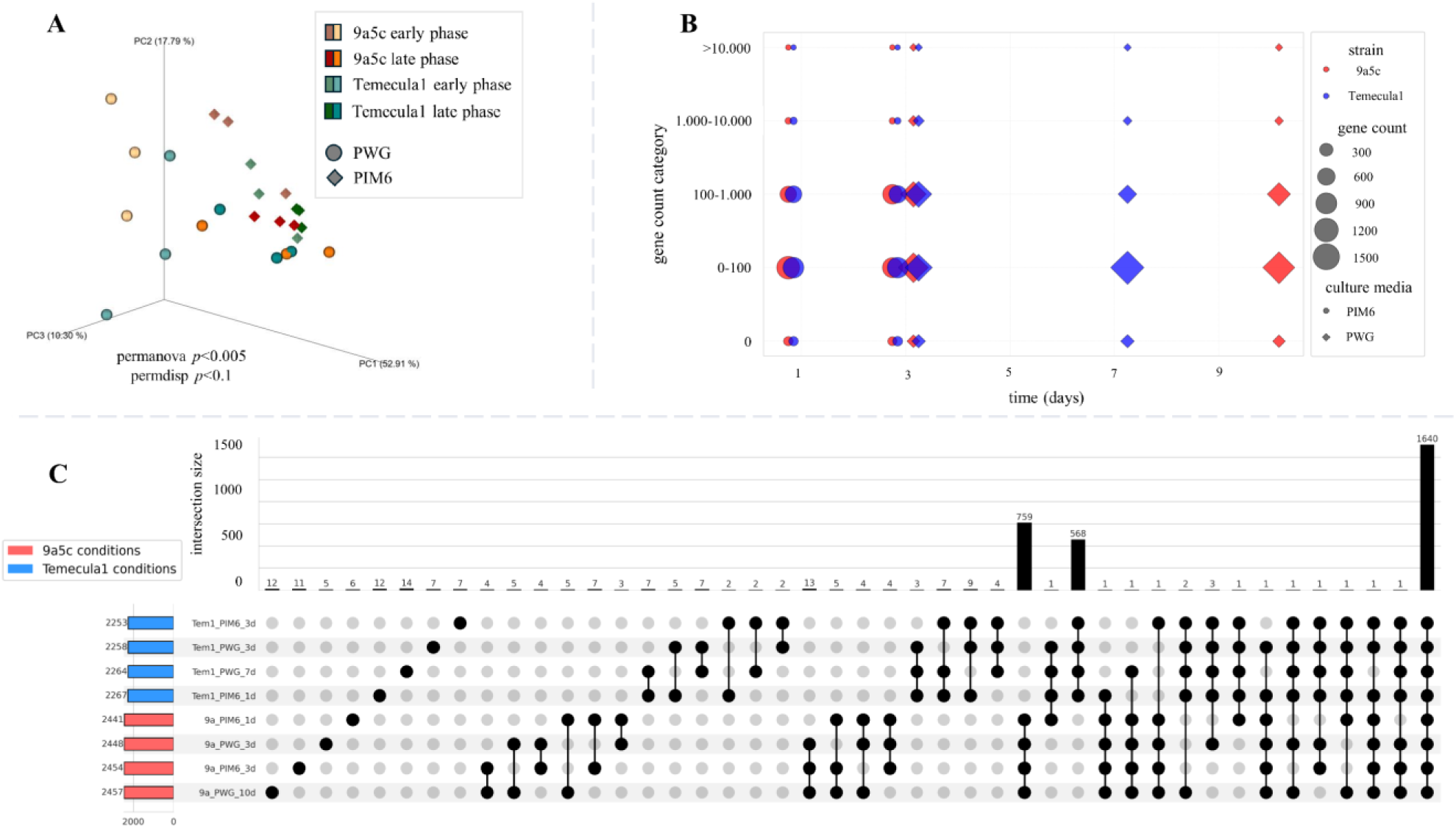
Global transcriptome across *X. fastidiosa* strains, media, and growth phases. **(A)** Principal Coordinates Analysis (PCoA) based on Bray–Curtis dissimilarity of normalized transcriptomic profiles. Samples cluster according to strain (9a5c vs. Temecula1), culture medium (PIM6 vs. PWG), and growth phase (early vs. late exponential). **(B)** Bubble plot summarizing Transcripts per Million (TPM) counts for genes in all pairwise comparisons between conditions. Each bubble represents a comparison, with size proportional to the number of TPMs, and **(C)** UpSet plot depicting the intersection of TPM sets across selected conditions. Horizontal bars show the total number of TPMs per comparison, and vertical bars represent the size of each intersection. Connected dots indicate shared TPMs across multiple contrasts, highlighting condition-specific and common transcriptional responses.

We also observed genes having expression values equal to zero in all transcriptomes (6.9% to 9.0%). Among them, the great majority encoded tRNAs and rRNAs, once their transcripts were depleted during the sequencing library preparation protocol. Other genes having no expression were mainly those with only a hypothetical function and those from prophage regions. This data suggests these genes belong to remnant regions of incomplete and inactive phages or that the culture conditions did not stimulate their expression. Despite most phage-related genes being silenced, a few were expressed. An UpSet analysis of TPM sets showed that 1,640 genes were expressed in every condition, with 759 genes shared across all four 9a5c transcriptomes and 568 genes shared across all four Temecula1 transcriptomes; all other intersections (or singletons) comprised fewer than 20 genes (Figure 3C).

Transcripts were also mapped to the plasmid sequences present in both strains (Marques et al., 2001; Simpson et al., 2000; Van Sluys et al., 2003). No expression of the two genes in the small plasmid present in both strains, pXF1.3, was seen in any condition tested. In contrast, all 69 predicted genes in plasmid pXF51 from strain 9a5c (Marques et al., 2001) were expressed in most of the transcriptomes. The only exceptions were two replicates in PIM6 during early exponential phase, where one gene of unknown function was not expressed (pXF51_00066). All of the expressed genes exhibited high TPM expression values in both media, commonly “above 10,000” and “between 10,000 and 1,000”. This high expression might reflect the presence of multiple plasmid copies. In plasmid pXF51 of strain 9a5c, the most abundant transcript was for a phage-related protein (pXF51_00036) and was observed in all transcriptomes. Fourteen operons were found in plasmid pXF51, most consisting of only two genes, similar to those predicted by the DOOR^2^ database (Table 1).

### Stress, competition, and metabolic readiness

Several of the most highly expressed CDSs in all transcriptomes encode bacteriocins and toxins, including the three colicins V (XF0262/PD0215, XF0263/PD0216, and XF0264/PD0217) described in the *X. fastidiosa* 9a5c genome (Simpson et al., 2000; Van Sluys et al., 2003) and newly identified microcins (XF1217/PD0497, XF1218/PD0498, and XF1306/-), previously annotated as hypothetical proteins but now supported by functional and expression data (Duarte and Da Silva, 2013). We also observed that the gene that encodes protein Ax21/Omp1X (XF1803/PD1063) was among those most highly expressed (Supplementary Tables S7-8).

The most abundant transcripts also include those for chaperone proteins GroEL and GroES, and also the RNA chaperone Hfq (Figure 4A). While these proteins are commonly abundant in bacterial cells, their high expression levels here may reflect increased cellular stress under *in vitro* culture conditions. Transcripts for bacterioferritin and peroxiredoxin were also quite abundant, reinforcing this hypothesis. Other genes with high expression encode cold shock proteins and the sigma factors σ^70^ and σ^32^ (RpoH), pilins (FimA), outer membrane proteins (such as OmpW), peptidases and proteases (such as Clp), among others. Interestingly, genes with hypothetical function, e.g., XF1024/PD0312, XF1217PD0497, and XF1287/PD0540, also showed high expression values in several transcriptomes, and thus, are good candidates for future studies to characterize their functions. A previously curated list of 73 well-characterized virulence genes (Uceda-Campos et al., 2022), categorized into sessile- and mobile-phase groups, was cross-referenced with our expression dataset (Figure 4B). In general, these were downregulated in late growth phases. In PIM6, their expression in Temecula1 resembles the pattern described by Chatterjee (2008) with mobility-related (plant colonization-phase) genes declining over time, while those contributing to a sessile phenotype (insect biofilm phase) genes become progressively upregulated.

**Figure 4.**
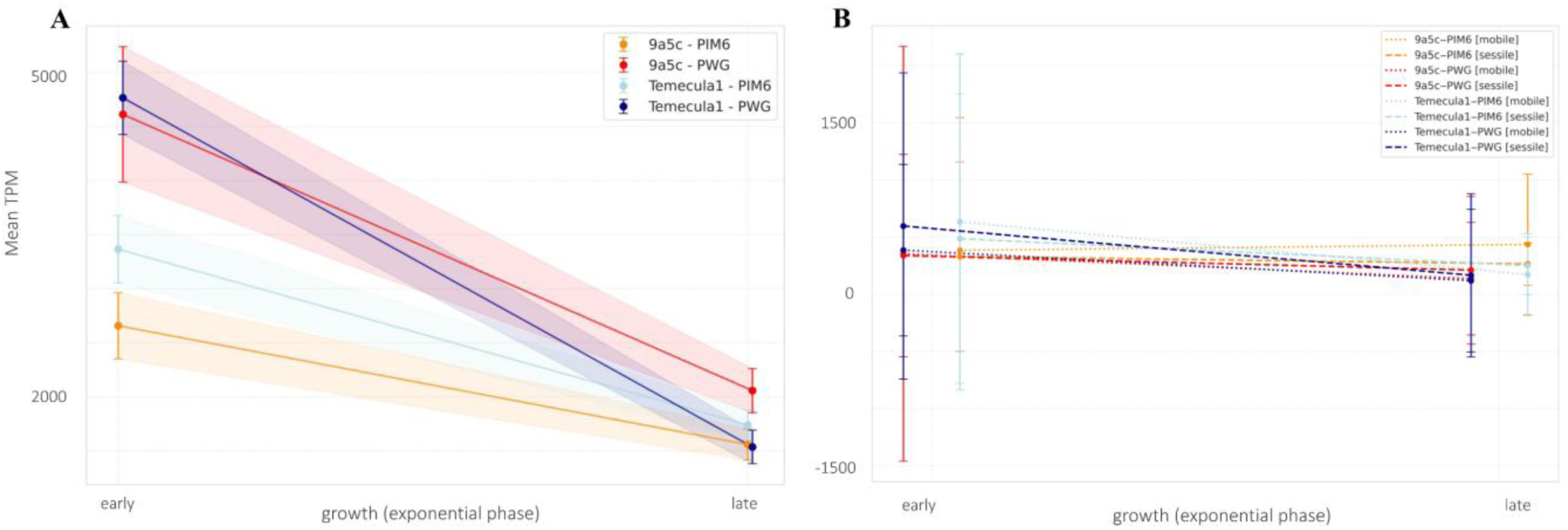
Expression trends of highly expressed and virulence-associated genes in *X. fastidiosa* strains. (**A**) Top 100 expressed shared genes. Line plots represent the mean TPM values (± standard error) of the 100 most highly expressed genes common to strains 9a5c and Temecula1, averaged across biological replicates for each condition. Conditions are organized by strain, medium (PIM6 vs. PWG), and growth phase (early vs. late). Statistical differences among strain– condition groups were assessed using one-way ANOVA followed by Benjamini–Hochberg FDR correction and Tukey’s HSD post hoc test. (**B**) TPM expression trends for sessile- and mobile-phase genes. Mean TPM per gene (± SE) across early and late exponential phases in PIM6 and PWG, for both 9a5c and Temecula1. Colors identify each strain-medium combination, while dashed lines trace sessile-phase gene groups and dotted lines trace mobile-phase gene groups.

Gene rankings based on gene-normalized expression data (TPM) were also used for a global analysis of which metabolic pathways were active under the tested conditions. All transcriptomes of strains 9a5c and Temecula1 were analyzed, considering genes as “expressed” if they had TPM expression values greater than 10. The results were tabulated with a binary notation of “1”, indicating the presence of the route in that transcriptome, and “0”, indicating its absence (Table S9).

All essential metabolic pathways for growth and proliferation were active, including glycolysis/gluconeogenesis, the TCA cycle, pentose phosphate and glyoxylate pathways, oxidative phosphorylation, and pyruvate and ubiquinone metabolism. Apparent metabolism of other carbohydrates such as starch, sucrose, fructose, and mannose was also detected, although no evidence of galactose metabolism was found.

All housekeeping pathways for maintenance and transmission of genetic information, such as DNA replication, transcription, translation, and aminoacyl-tRNA synthesis, were active in all transcriptomes. In addition, DNA repair systems using base or nucleotide excision and mismatch repair were active. Genes involved in homologous recombination were also expressed. The pathways for purine and pyrimidine metabolism were also found in both strains. However, no evidence of sugar nucleotide metabolism for the synthesis of glycoconjugates was seen.

The pathways for protein export and types II and IV secretion systems were active in all transcriptomes. Other transport routes were also observed, such as ABC transporters, two-component systems, and sugar phosphotransfer system (PTS).

Pathways related to membrane and cell wall synthesis, such as glycerophospholipid metabolism, peptidoglycan and lipopolysaccharide biosynthesis, were all active, as well as various routes for saturated and unsaturated fatty acid and steroid biosynthesis in both strains. Notably, however, fatty acid and glycerolipid metabolism were only active in strain 9a5c, but not in Temecula1.

Several vitamins and coenzymes, such as thiamine, riboflavin, vitamin B6, biotin, porphyrin, lipoic acid, nicotinate and nicotinamide, pantothenate and coenzyme A, were also represented in the transcriptomes. However, ascorbate (vitamin C) metabolism was inactive in Temecula1 but active in 9a5c, and inositol metabolism was not found to be active in either strain.

Interestingly, MinPath-based functional inference indicated the presence of biosynthetic routes associated with antibiotics and polyketides, including pathways annotated for vancomycin, ansamycin, tetracycline, and streptomycin. While these findings suggest conserved metabolic capabilities, they do not necessarily imply active antibiotic production by *X. fastidiosa*.

Pathways for nitrogen, sulfur, glutathione metabolism, and processing of amino groups via the urea cycle were active. However, the absence of several amino acid biosynthetic pathways may contribute to the slow growth of *X. fastidiosa* and its reliance on supplemented media. Biosynthesis and degradation routes for alanine, aspartate, glycine, serine, threonine, arginine, proline, histidine, phenylalanine, and methionine were active. However, no evidence was found for glutamate, cysteine, tyrosine, glutamine and glutamate, selenium, and cyanamino acids catabolism. For lysine, valine, leucine, and isoleucine, only evidence for their biosynthesis, but not catabolism, was seen. Tryptophan metabolism was detected only in strain 9a5c, specifically in a subset of early- and late-phase replicates.

Interestingly, methane metabolism was inactive in these same transcriptomes but active in all others, suggesting a potential, albeit speculative, inverse relationship between these pathways. Conversely, methane metabolism was active in all other transcriptomes. Although this opposing pattern is intriguing, any potential biological relationship between the two pathways remains speculative.

In summary, metabolic pathway activity was largely conserved between strains 9a5c and Temecula1, with only minor differences observed. All essential routes for growth and survival were active across media and growth phases, indicating a high degree of adaptability.

### Differential expression and functional enrichment

We performed three types of comparisons: (A) within each strain and medium, early vs. late growth; (B) within each strain, PIM6 vs. PWG at matching growth phases; and (C) between strains under the same medium and growth phase. Table 2 summarizes the numbers of differentially expressed genes (DEGs; p ≤ 0.05; |log2FC| > 1) for each comparison. Overall, PIM6 elicited more transcriptional changes than PWG (especially during early exponential growth), and the largest between-strain differences were observed at early/mid phases.

**Table 2.** Summary of DEGs (p≤0.05; log2FC> |1|) in all comparisons.

| Level of comparison | Total genes | Condition 1 (c1) | Condition 2 (c2) | DEG* ($p \leq 0.05$; $\log_2FC > 1.0$) | Up-regulated genes (c1/c2) |
| --- | --- | --- | --- | --- | --- |
| **A** | 2,639 | 9a5c_PIM6_1d | 9a5c_PIM6_3d | 83 (3.1%) | 79/4 |
|  | 2,639 | 9a5c_PWG_3d | 9a5c_PWG_10d | 0 | 0 |
|  | 2,476 | Tem1_PIM6_1d | Tem1_PIM6_3d | 61 (2.5%) | 50/11 |
|  | 2,476 | Tem1_PWG_3d | Tem1_PWG_7d | 1 (0.04%) | 0/1 |
| **B** | 2,639 | 9a5c_PIM6_1d | 9a5c_PWG_3d | 4 (0.15%) | 0/4 |
|  | 2,639 | 9a5c_PIM6_3d | 9a5c_PWG_10d | 20 (0.8%) | 3/17 |
|  | 2,476 | Tem1_PIM6_1d | Tem1_PWG_3d | 1 (0.04%) | 1/0 |
|  | 2,476 | Tem1_PIM6_3d | Tem1_PWG_7d | 6 (0.24%) | 2/4 |
| **C** | 1,882 | 9a5c_PIM6_1d | Tem1_PIM6_1d | 102 (5.4%) | 54/48 |
|  | 1,882 | 9a5c_PIM6_3d | Tem1_PIM6_3d | 238 (12.6%) | 115/123 |
|  | 1,882 | 9a5c_PWG_3d | Tem1_PWG_3d | 58 (3.1%) | 42/16 |
|  | 1,882 | 9a5c_PWG_10d | Tem1_PWG_7d | 95 (5.0%) | 46/49 |

\*DEG: differentially expressed genes; Tem1: Temecula1.

To gain a systems-level understanding of the transcriptomic changes, we performed GO (Ashburner et al., 2000) enrichment (padj ≤ 0.05; |FC| > 2) with BayGO (Vêncio et al., 2006) for each comparison. In PIM6, early vs. late exponential phases showed enrichment of shared categories (protein biosynthesis, ribosome biogenesis, RNA polymerase activity) in both strains (Figure 5A; Table S10).

**Figure 5.**
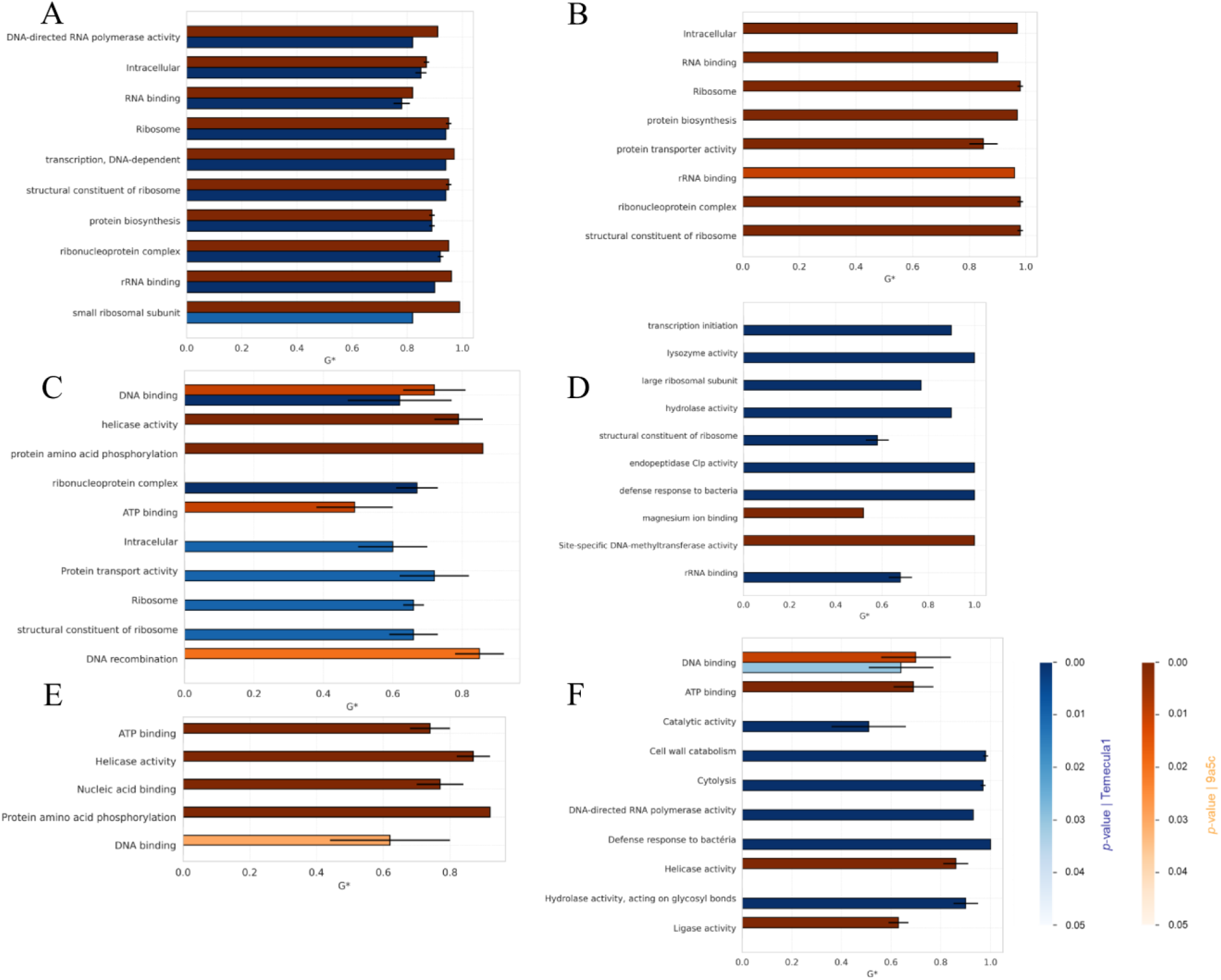
Comparative gene ontology (GO) enrichment analysis across growth phases, media, and *X. fastidiosa* strains. Top 10 significantly enriched GO terms (biological process category) shared between strains 9a5c and Temecula1, based on adjusted P-values ≤ 0.05. Each plot compares normalized enrichment scores (G\*) across different experimental conditions, with orange-shade bars representing 9a5c and blue-shade bars representing Temecula1. GO terms are ordered by increasing mean P-value across strains. In conditions where one strain lacked enriched GO terms (e.g., Temecula1 under certain comparisons), G\* values were set to zero for visualization purposes to preserve consistency in shared term comparisons. **A.** Comparison between early and late exponential phases in PIM6 medium for both strains. **B.** Comparison between late exponential growth in PWG (rich medium) and PIM6 (minimal medium) for strain 9a5c. **C.** Cross-strain comparison (9a5c vs Temecula1) during early exponential phase in PIM6 medium. **D.** Cross-strain comparison during late exponential phase in PIM6 medium. **E.** Cross-strain comparison during early exponential phase in PWG medium. **F.** Cross-strain comparison during late exponential phase in PWG medium. G\* represents a normalized enrichment score accounting for gene set size and background, enabling comparisons between conditions. Terms shown were selected based on overlap across strains and are visualized with up to 10 GO terms per panel. Full enrichment tables are provided in Supplementary Tables S10–S15.

For 9a5c, late PWG versus late PIM6 revealed enrichment exclusively in protein synthesis and transport pathways, including ribosome assembly (Figure 5B; Table S11). No enriched terms were detected for Temecula1 in this comparison. In the early-phase cross-strain comparison in PIM6, 9a5c was enriched in DNA ligation/recombination, helicase, and ligase activities, whereas Temecula1 was enriched in protein synthesis and transport (Figure 5C; Table S12). In late-phase PIM6, 9a5c showed enrichment for methyltransferase activity and DNA methylation. Conversely, Temecula1 exhibited extensive enrichment (28 terms), spanning transcription, ATP metabolism, defense responses (lysozyme, cytolysis, peptidoglycan catabolism), and protein synthesis (Figure 5D; Table S13), suggesting heightened metabolic adaptation to nutritional stress. In PWG, 9a5c maintained DNA-ligation and amino-acid phosphorylation enrichment in both phases, while Temecula1 in the late phase exhibited enrichment in carbohydrate metabolism, transcription, and defense (Figures 5E-F; Tables S14-15).

We next focused on known *X. fastidiosa* virulence genes (Chatterjee et al., 2008; Simpson et al., 2000; Van Sluys et al., 2003), as presented in Figure 6 (Table S16). In PIM6, DEGs appeared only during early exponential growth: both strains up-regulated *pilV* (twitching motility) (Meng *et al*., 2005), *fimA* (adhesion) (Li et al., 2007), and a zinc-dependent chaperone protease; 9a5c alone up-regulated *pilO* and *pilQ*. In late-phase media comparisons, 9a5c up-regulated *pilQ* in PWG, while Temecula1 up-regulated a σ54-modulating regulatory protein in PIM6 (Ancona et al., 2014; Oosthuizen et al., 2002). In early-phase cross-strain PIM6, five virulence genes were differentially expressed: 9a5c up-regulated PilY1, a histidine kinase, and an AraC-type regulator; Temecula1 up-regulated a pre-pilin peptidase and a lipase—patterns maintained in late PIM6. In late PIM6, 9a5c up-regulated a glycosidase, a LuxR/UhpA family regulator, flavodoxin WrbA, another transcriptional regulator, and a hemolysin; Temecula1 up-regulated *rpfG/F* (quorum sensing), σ70/RpoH, a nitrogen regulator, ClpP, a zinc protease, and a PHA-associated protein. In PWG, 9a5c consistently up-regulated PilY1 and a histidine kinase in both phases, plus a hemolysin early; Temecula1 up-regulated only a lipase in late phase.

**Figure 6.**
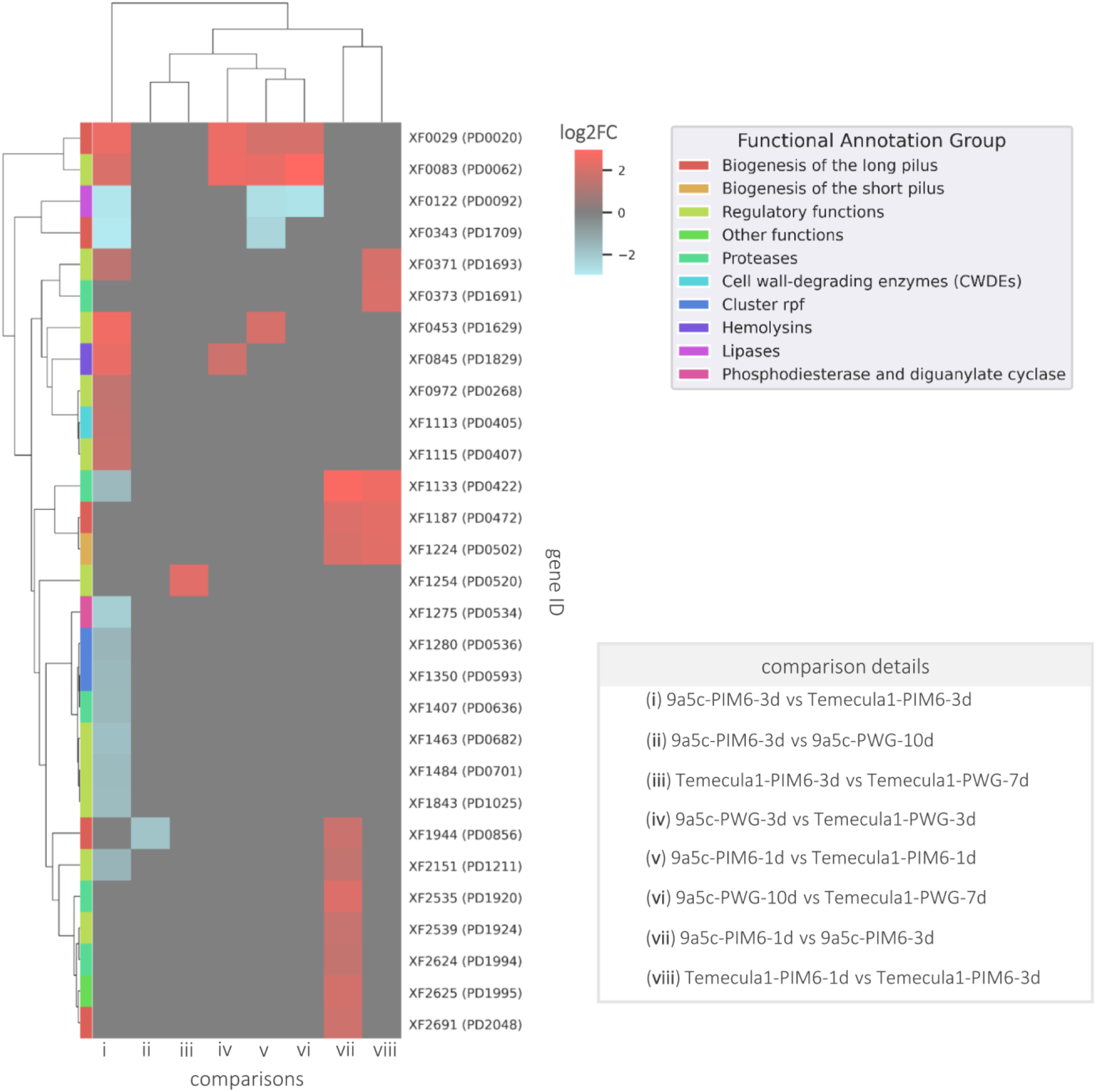
Clustermap of differentially expressed genes (DEGs) across experimental comparisons, grouped by functional annotation. Each column represents a DEG, colored according to its assigned functional annotation group (legend, bottom right). Rows correspond to pairwise transcriptomic comparisons between conditions, including distinct media (PIM6 vs. PWG), time points (early vs. late exponential phases), and strain contrasts (9a5c vs. Temecula1). The heatmap displays log₂ fold-change values, where red indicates upregulation, blue indicates downregulation, and grey denotes no change (or gene not differentially expressed). Hierarchical clustering was applied to both genes and comparisons based on fold-change patterns. Only DEGs shared across comparisons were included.

## DISCUSSION

Our RNA-Seq atlas of *X. fastidiosa* strains Temecula1 and 9a5c reveals a transcriptional landscape finely linked to both ecological niches and genomic lineage. By profiling gene expression across xylem-mimicking minimal medium (PIM6) and nutrient-rich medium (PWG) at early and late exponential phases of growth, we uncovered a multifaceted regulatory plasticity—from operon shifts to the activity of non-coding RNAs—that supports *X. fastidiosa*’s remarkable adaptability to the distinct niches it occupies. Transcriptional differences between strains often outweighed medium-induced changes, underscoring the importance of intra-species diversity when dissecting virulence strategies or designing control interventions.

Phenotypic and growth-curve assays provided crucial context. In PIM6, both strains transitioned into the stationary phase by day 3, whereas the richer PWG extended exponential growth beyond 10 days, especially in 9a5c. These kinetics align with prior findings in sap-like media (Bellato et al., 2004; Zaini et al., 2009) but diverge from the slower or irregular growth observed in other defined culture media (Ciraulo et al., 2010). Temecula1’s delayed growth in PWG relative to 9a5c likely reflects its heightened nutritional demands and perhaps a more copiotrophic lifestyle —an observation with practical implications for culturing diverse field isolates and interpreting *in vitro* assays of biofilm formation or virulence potential.

At the level of transcriptional organization, we detected a higher number of expressed operons in 9a5c. While partly proportional to genome size, these numbers also reflect regulatory divergence. Comparisons with operons expressed in Temecula1 under different conditions (Parker et al., 2016) revealed only 318 out of 386 overlapping units, highlighting widespread conditional operon reconfiguration. This operon plasticity likely enables *X. fastidiosa* to rapidly remodel transcriptional units in response to subtle shifts in host-derived signals or nutrient availability, allowing modular deployment of metabolic and virulence cassettes.

Two small RNAs were highly expressed in all conditions: the RNA component of RNase P (*rnpB*) and 6S RNA. *rnpB* dominated the expression landscape, consistent with its critical role in tRNA maturation and global RNA processing (Guerrier-Takada et al., 1983). Meanwhile, 6S RNA, a universal transcriptional regulator that binds and sequesters σ⁷⁰-RNAP complexes, had a high level of expression in PIM6 and during stationary phase. This suggests that *X. fastidiosa* uses 6S RNA to modulate global transcription in response to nutrient limitation, mirroring strategies observed in *E. coli* and other bacteria facing stationary-phase stress (Steuten et al., 2014; Wassarman and Storz, 2000).

We observed robust expression of bacteriocins (including colicin- and microcin-like proteins) and the Ax21/Omp1X porin across all conditions. Despite growth in monoculture, these systems indicate an intrinsic readiness to compete within polymicrobial xylem niches – perhaps most likely within insect vectors. Additionally, a RelE-like toxin encoded on plasmid pXF51 ranked among the highest plasmid transcripts, linking toxin–antitoxin (TA) systems to stress adaptation and persister cell formation (Burbank and Stenger, 2017; Harms et al., 2018). These findings reinforce the view that *X. fastidiosa* maintains latent but readily deployable tools for niche domination and long-term survival in both plant and insect hosts. Differential gene expression analysis revealed that PIM6 triggered approximately fivefold more changes in gene expression than PWG, consistent with a broad transcriptional maintenance under nutrient stress. However, overlap of DEGs across matched growth phases in PIM6 was limited. This suggests that *X. fastidiosa* maintains a “poised” transcriptional program—ready for rapid adaptation but avoiding unnecessary reprogramming. Conversely, strain-specific comparisons showed that up to 12.6% of orthologs were differentially expressed, often with fold-changes far exceeding those seen in media-dependent contrasts. This emphasizes the significant transcriptional divergence among strains and advises caution against overgeneralization from single-strain studies.

GO enrichment analyses highlighted strain-specific stress responses. In PIM6, Temecula1 upregulated genes associated with transcriptional regulation, ATP metabolism, lysozyme activity, and ribosome biogenesis—hallmarks of a robust stress adaptation strategy. In contrast, 9a5c favored DNA ligation, helicase activity, and methylation processes, potentially reflecting enhanced genomic plasticity under xylem-mimicking stress. In PWG, both strains reduced these expressions, although Temecula1 retained moderate defense-related responses during late growth, suggesting a constitutively cautious transcriptional phenotype.

Virulence-linked expression patterns followed a similar pattern. In early PIM6 growth, both strains upregulated long-pilus assembly genes (*pilV*, *pilO*, *pilQ*) and the short-pilus adhesin *fimA*, potentially priming for motility and host colonization. Later on, divergent strategies emerged: Temecula1 activated quorum-sensing genes (*rpfF*, *rpfG*), the lipase PD1211, and multiple sigma factors, while 9a5c favored glycosidase enzymes and LuxR-family regulators. The near-absence of virulence-related differential expression in PWG suggests that rich medium conditions suppress the “invasion-ready” state—an insight with clear implications for interpreting *in vitro* pathogenicity assays.

Altogether, our findings describe *X. fastidiosa* as a “transcriptionally alert pathogen”, maintaining virulence and stress-response systems in a standby mode while fine-tuning operon use, small RNA activity, and toxin expression in response to environmental cues. From a disease control standpoint, this adaptability highlights regulatory constrictions—such as 6S RNA, TA modules, and type IV pilus components—as promising targets for broad-spectrum attenuation. Future studies incorporating *in planta* time-course RNA-Seq, single-cell resolution, and functional validation of alternative transcription start sites will further inform real-time regulatory dynamics underlying *X. fastidiosa*’s persistent colonization of xylem vessels and insect vectors.

## AUTHORS CONTRIBUTIONS

PMP and AMDS contributed to the conception and design of the study; PMP performed the experiments; PMP and ORF-J performed the data collection; PMP and JMJ performed sequence mapping; PMP and ORF-J performed the statistical analysis; DB performed metabolic pathway analysis; PP, ORF-J, and PAZ contributed to the discussion of results; PMP, ORF-J, PAZ, and AMDS wrote the manuscript. All authors contributed to the manuscript revision, read and approved the submitted version.

## ACKNOWLEDGEMENTS

We thank: Prof. João Carlos Setubal (computational resources); Prof. Steven E. Lindow (Temecula1 strain and scientific insights); and Prof. Michele Igo (PIM6 medium protocol). This work honors the enduring legacy of Prof. Aline Maria da Silva, whose mentorship was foundational to our research.

## CONFLICT OF INTEREST

The authors declare that the research was conducted in the absence of any commercial or financial relationships that could be construed as a potential conflict of interest.

## FUNDING

This research was funded by the São Paulo Research Foundation (FAPESP), grant number 08/11703-4, and by the National Council for Scientific and Technological Development (CNPq), grant number 476656/2012-5, as well as the Coordination for the Improvement of Higher Education Personnel (CAPES), grant number 3385/2013. Fellowships from CAPES supported ORF-J, JMJ, and DB. The FAPESP fellowship 2011/24091-0 supported PMP. A.M.d.S. received a research fellowship award (309182/2016-6) from the National Council for Scientific and Technological Development (CNPq).

## DATA AVAILABILITY

Raw RNA reads from this study will be publicly available in the GenBank/NCBI Sequence Read Archive (SRA) before peer-review submission.

## SUPPLEMENTARY MATERIAL

The following supporting information is available separately: Table S1. Summary of the yield of each sequenced library; Table S2. Pearson Correlation table of all-against-all conditions and their replicates; Table S3. List of untranslated regions (UTRs) obtained from sequence mapping of the transcriptomes of strain 9a5c and their respective coordinates. Mapping was performed using Rockhopper2 software; Table S4. List of untranslated regions (UTRs) obtained from sequence mapping of the transcriptomes of the Temecula1 strain and their respective coordinates. Mapping was performed with Rockhopper2. software; Table S5. List of operons predicted from RNA-Seq data of strain 9a5c. Prediction was performed with Rockhopper2 software; Table S6. List of operons predicted from RNA-Seq data of the Temecula1 strain. The prediction was performed with the Rockhopper2 software; Table S7. List of genes transcribed from strain 9a5c, based on TPM values; Table S8. List of genes transcribed from strain Temecula1, based on TPM values; Table S9. Analysis of metabolic pathways present (1) or absent (0) in the transcriptomes of strains 9a5c and Temecula1; Table S10. Comparison between early and late exponential phases in PIM6 medium for both strains; Table S11. Comparison between late exponential growth in PWG (rich medium) and PIM6 (minimal medium) for strain 9a5c; Table S12. Cross-strain comparison (9a5c vs Temecula1) during early exponential phase in PIM6 medium; Table S13. Cross-strain comparison during late exponential phase in PIM6 medium; Table S14. Cross-strain comparison during the early exponential phase in PWG medium; Table S15. Cross-strain comparison during late exponential phase in PWG medium; Table S16. lists of identities of differentially expressed virulence-associated genes identified within each comparison.

## Notes

### Competing Interest Statement

The authors have declared no competing interest.

